# TORC1 inhibition as an immunotherapy to reduce infections in the elderly

**DOI:** 10.1101/230813

**Authors:** Joan B. Mannick, Melody Morris, Hans-Ulrich P. Hockey, Guglielmo Roma, Martin Beibel, Kenneth Kulmatycki, Mollie Watkins, Tea Shavlakadze, Weihua Zhou, Dean Quinn, David J. Glass, Lloyd B. Klickstein

**Affiliations:** Novartis Institutes for Biomedical Research, 181 Massachusetts Avenue, Cambridge, MA 02139, USA; Biometrics Matters Limited, 13 Nevada Rd, Hamilton 3216 New Zealand; Novartis Institutes for Biomedical Research, Fabrikstrasse-22, Novartis Campus, 4056 Basel, Switzerland; P3 Research, PO Box 7366, Newtown Wellington 6242 New Zealand

## Abstract

mTOR inhibition extends lifespan and ameliorates aging-related pathologies including declining immune function in model organisms. The objective of this Phase 2a clinical trial was to determine if low dose mTOR inhibitor therapy enhanced immune function and thereby decreased infection rates in elderly subjects. The results indicate that 6 weeks of treatment with a low dose combination of a catalytic (BEZ235) plus an allosteric (RAD001) mTOR inhibitor (that selectively inhibits TORC1 downstream of mTOR) was safe, significantly decreased the rate of infections reported by elderly subjects for a year following study drug initiation, upregulated antiviral gene expression, and significantly improved influenza vaccination response. Thus selective TORC1 inhibition with a combination of BEZ235 and RAD001 may be efficacious as immunotherapy to reduce infections, a leading cause of death in the elderly.

**One Sentence Summary:** Treatment of elderly subjects with a low dose mTOR inhibitor regimen that selectively inhibits TORC1 significantly decreased infection rates

---

Aging may be due, in part, to alterations of a discrete set of cell signaling pathways including the mTOR pathway (*1*). Inhibition of the mTOR pathway has extended lifespan in every species studied to date suggesting that mTOR is an evolutionarily conserved pathway that may regulate aging (*2*). One of the aging-related conditions that improves in old mice treated with mTOR inhibitors is immunosenescence (the decline in immune function that occurs during aging) (*3*). Immunosenescence leads to increased rates of infections including respiratory tract infections that are the 4rth leading cause of death in people ≥ 85 years of age, and the 8^th^ leading cause of death in people ≥ 65 years of age in the United States (*4,5*). Therefore, infection-related morbidity and mortality in the elderly may be substantially reduced if mTOR inhibition enhances the ability of the aging immune system to fight infectious pathogens.

mTOR signals via two complexes: TORC1 and TORC2. Many of the beneficial effects of mTOR inhibition on aging may be mediated by inhibition of TORC1 (*6,7*). In contrast, TORC2 inhibition has been associated with adverse events including hyperglycemia and hypercholesterolemia, and with decreased lifespan in male mice (*7,8*). In addition, several long-lived animal models have increased rather than decreased TORC2 activity (*9*). Therefore, optimal mTOR inhibition for the treatment of aging-related conditions such as immunosenescence may be a regimen that inhibits TORC1 without inhibiting TORC2.

Rapalogs such as RAD001 are a class of allosteric mTOR inhibitors that consistently inhibit only S6K downstream of TORC1 (*10*). BEZ235 is a dual PI3K/mTOR ATP-competitive catalytic site inhibitor that at high concentrations inhibits TORC1, TORC2 and PI3K, but at low concentrations (≤ 20 nM) mainly inhibits S6K phosphorylation and to a lesser extent 4EPB1 phosphorylation downstream of TORC1 (*11, 12, 13*). However, low doses of BEZ235 (and other catalytic mTOR inhibitors) in combination with low doses of RAD001 synergistically inhibit multiple nodes downstream of TORC1 without inhibiting TORC2 activity (*14,15*). Thus the combination of a low dose rapalog plus a catalytic site mTOR inhibitor may be more efficacious than monotherapy with a rapalog or catalytic site mTOR inhibitor for the treatment of aging-related conditions such as immunosenescence. In a previous study we demonstrated that treatment with the rapalog RAD001 improved immune function in elderly volunteers as assessed by response to influenza vaccination (*16*). In the current study we extended these findings and assessed whether 1) a low dose combination of a rapalog (RAD001) plus a catalytic site mTOR inhibitor (BEZ235) resulted in a greater improvement in immune function in elderly subjects than monotherapy with either RAD001 or BEZ235; and 2) whether low dose mTOR inhibitor treatment resulted in sufficient immune enhancement to not only improve influenza vaccination response, but also to decrease total infection rates in the elderly at doses that were safe.

A total of 264 elderly volunteers ≥ 65 years of age, without unstable medical conditions, were enrolled in a randomized, double-blinded, placebo-controlled trial at 12 clinical sites (Fig. 1). Subjects were assigned randomly to receive one of four oral mTOR inhibitor dosing regimens or a corresponding matching placebo: RAD001 0.5 mg once daily, RAD001 0.1 mg once daily, BEZ235 10 mg once daily, or a combination of 0.1 mg RAD001 and 10 mg BEZ235 once daily. The placebo groups were pooled for analysis.

**Fig. 1.**
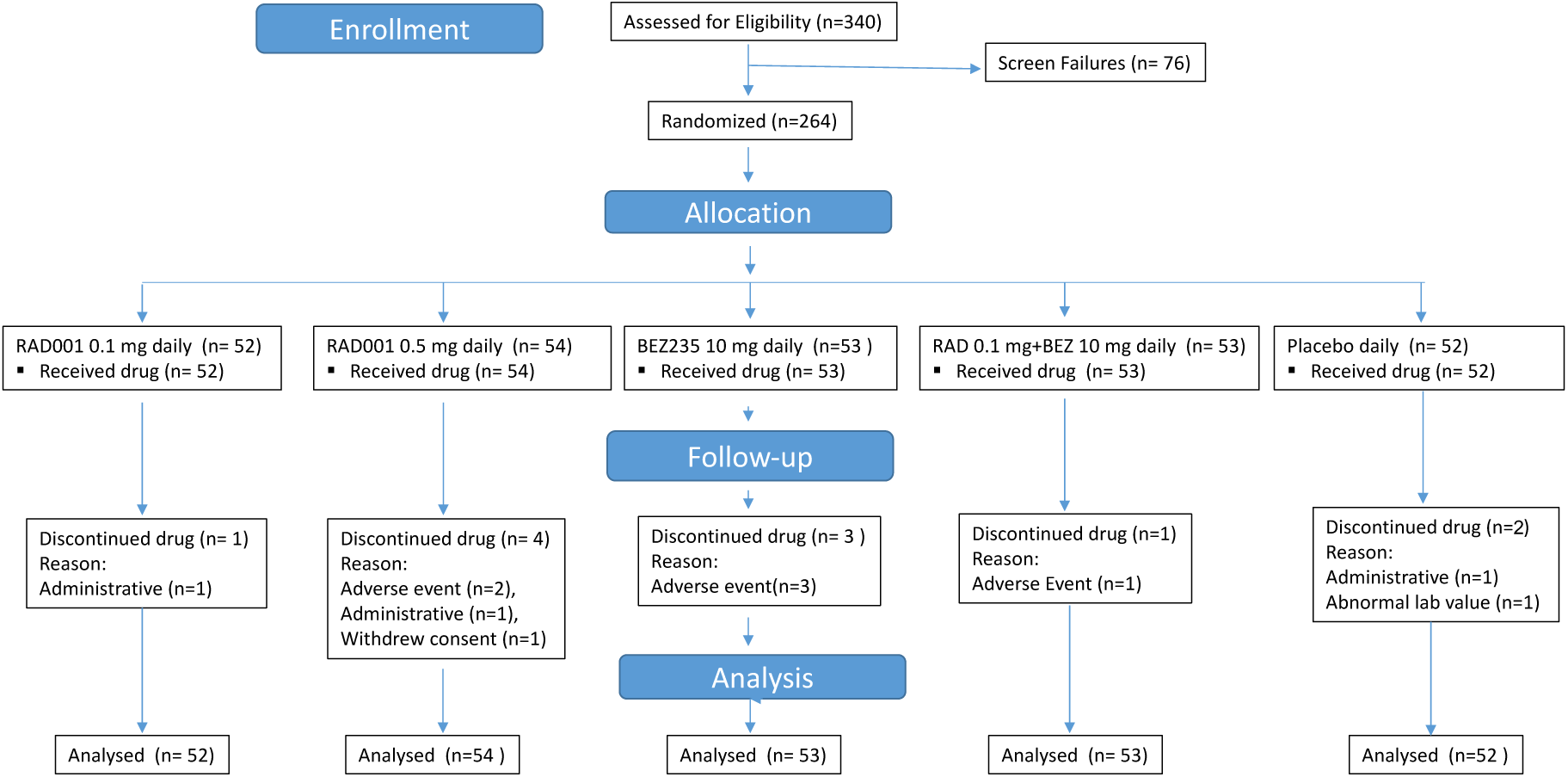
Study Diagram.

Subjects were treated for 6 weeks with study drug and after a 2 week drug-free interval, were given a seasonal influenza vaccine (Fluvax®, CSL Biotherapies). Antibody titers to the 3 strains of influenza in the vaccine (A/H1N1, A/H2N3 and B) were measured in serum collected at baseline (after 6 weeks of study drug administration but prior to vaccination) and 4 weeks after vaccination. The subjects were then followed for approximately 9 months off study drug (approximately one year following initiation of study drug treatment).

Baseline demographics between the treatment arms were similar (Supplementary Table 1). Of the 264 subjects enrolled, 253 completed the study (Fig. 1). In general, the mTOR inhibitor regimens were well tolerated. No deaths occurred during the study. 26/264 participants experienced at least one serious adverse event (SAE) during the 12 months they were followed in the study. There were no significant differences in the percentage of subjects experiencing SAEs between treatment groups and placebo: 9.6 % in the 0.1 mg RAD001 daily cohort, 13% in the 0.5 mg RAD001 daily cohort, 9.4% in the BEZ235 10 mg daily cohort, 7.5% in the 0.1 mg RAD001 + 10 mg BEZ235 cohort, and 9.6% in the placebo cohort. Only one SAE (syncope in a subject in the placebo cohort) was deemed by an investigator to be related to study drug. A list of adverse events (AEs) that occurred in more than 5% of subjects in any treatment group in the safety population (defined as any subject who received at least one partial dose of study drug) is provided in Table 1. Diarrhea was the most frequently reported adverse event that occurred more often in all the mTOR inhibitor cohorts than in the placebo treatment group and was of mild severity in the majority of cases. Of note, rates of hyperglycemia and hypercholesterolemia (adverse events associated with TORC2 inhibition) were lower in the mTOR inhibitor treatment groups than the placebo treatment group suggesting that the mTOR inhibitor treatment regimens were not inhibiting TORC2.

**Table 1.** Incidence of AEs by preferred term > 5% incidence in any group during the 1 year study

|  | **RAD001**<br>**0.1 mg** | **RAD001**<br>**0.5 mg** | **BEZ235**<br>**10 mg** | **RAD001 0.1**<br>**mg and**<br>**BEZ235**<br>**10 mg** | **Placebo,**<br>**pooled** | **Total** |
| --- | --- | --- | --- | --- | --- | --- |
|  | **N=52**<br>**n (%)** | **N=54**<br>**n (%)** | **N=53**<br>**n (%)** | **N=53**<br>**n (%)** | **N=52**<br>**n (%)** | **N=264**<br>**n (%)** |
| Total AE(s) | 320 | 406 | 346 | 319 | 401 | 1792 |
| Patients with AE(s) | 50 (96.2) | 52 (96.3) | 52 (98.1) | 51 (96.2) | 51 (98.1) | 256 (97.0) |
| Upper respiratory tract infection | 11 (21.2) | 12 (22.2) | 11 (20.8) | 14 (26.4) | 16 (30.8) | 64 (24.2) |
| Headache | 7 (13.5) | 16 (29.6) | 13 (24.5) | 11 (20.8) | 9 (17.3) | 56 (21.2) |
| Nasopharyngitis | 14 (26.9) | 13 (24.1) | 6 (11.3) | 7 (13.2) | 10 (19.2) | 50 (18.9) |
| Diarrhoea | 10 (19.2) | 12 (22.2) | 10 (18.9) | 9 (17.0) | 4 (7.7) | 45 (17.0) |
| Cough | 7 (13.5) | 7 (13.0) | 6 (11.3) | 5 (9.4) | 10 (19.2) | 35 (13.3) |
| Fatigue | 6 (11.5) | 4 (7.4) | 8 (15.1) | 4 (7.5) | 6 (11.5) | 28 (10.6) |
| Oropharyngeal pain | 8 (15.4) | 2 (3.7) | 8 (15.1) | 1 (1.9) | 7 (13.5) | 26 (9.8) |
| Arthralgia | 4 (7.7) | 7 (13.0) | 6 (11.3) | 3 (5.7) | 5 (9.6) | 25 (9.5) |
| Fall | 4 (7.7) | 3 (5.6) | 6 (11.3) | 7 (13.2) | 5 (9.6) | 25 (9.5) |
| Nausea | 4 (7.7) | 6 (11.1) | 4 (7.5) | 7 (13.2) | 4 (7.7) | 25 (9.5) |
| Rhinitis | 3 (5.8) | 4 (7.4) | 3 (5.7) | 9 (17.0) | 6 (11.5) | 25 (9.5) |
| Dizziness | 5 (9.6) | 5 (9.3) | 6 (11.3) | 3 (5.7) | 4 (7.7) | 23 (8.7) |
| Back pain | 5 (9.6) | 0 | 8 (15.1) | 2 (3.8) | 5 (9.6) | 20 (7.6) |
| Contusion | 5 (9.6) | 2 (3.7) | 2 (3.8) | 7 (13.2) | 4 (7.7) | 20 (7.6) |
| Oral herpes | 5 (9.6) | 2 (3.7) | 7 (13.2) | 3 (5.7) | 3 (5.8) | 20 (7.6) |
| Urinary tract infection | 5 (9.6) | 5 (9.3) | 2 (3.8) | 0 | 7 (13.5) | 19 (7.2) |
| Lower respiratory tract infection | 3 (5.8) | 3 (5.6) | 1 (1.9) | 5 (9.4) | 6 (11.5) | 18 (6.8) |
| Gastroenteritis | 3 (5.8) | 4 (7.4) | 3 (5.7) | 5 (9.4) | 2 (3.8) | 17 (6.4) |
| Mouth ulceration | 5 (9.6) | 2 (3.7) | 2 (3.8) | 6 (11.3) | 2 (3.8) | 17 (6.4) |
| Muscle strain | 3 (5.8) | 4 (7.4) | 2 (3.8) | 3 (5.7) | 5 (9.6) | 17 (6.4) |
| Constipation | 3 (5.8) | 6 (11.1) | 4 (7.5) | 1 (1.9) | 2 (3.8) | 16 (6.1) |
| Bronchitis | 3 (5.8) | 3 (5.6) | 2 (3.8) | 3 (5.7) | 3 (5.8) | 14 (5.3) |
| Laceration | 4 (7.7) | 2 (3.7) | 2 (3.8) | 3 (5.7) | 3 (5.8) | 14 (5.3) |
| Musculoskeletal pain | 1 (1.9) | 8 (14.8) | 1 (1.9) | 4 (7.5) | 0 | 14 (5.3) |
| Pain in extremity | 1 (1.9) | 3 (5.6) | 3 (5.7) | 2 (3.8) | 5 (9.6) | 14 (5.3) |

Ordered by total frequency
Under one treatment, a participant with multiple occurrences of an adverse event is counted only once in the AE category

The ability of RAD001 and/or BEZ235 to improve immune function in elderly volunteers was evaluated by measuring the serologic response to the 2014 seasonal influenza vaccine. The primary endpoint of the study was a 1.2 fold increase relative to placebo in the hemagglutination inhibition (HI) geometric mean titer (GMT) ratio (GMT 4 weeks post vaccination/GMT at baseline) for at least 2 out of 3 influenza vaccine strains. This endpoint was chosen because the approximately 1.2-fold increase in the influenza GMT ratio induced by the MF-59 vaccine adjuvant was associated with a decrease in influenza illness (*17*).

In the modified intent-to-treat (ITT) population (those subjects who received at least one full dose of study drug), only the combination low dose RAD001 (0.1 mg daily) + BEZ235 (10 mg daily) met the primary endpoint of the study and resulted in a statistically significant greater than 20% increase in the influenza GMT ratio for 3/3 influenza vaccine strains (Fig. 2). RAD001 monotherapy (0.1 mg or 0.5 mg daily) resulted in a statistically significant greater than 20% increase in influenza GMT ratio for 1/3 influenza vaccine strains. BEZ235 monotherapy did not result in an increase in influenza GMT ratios for any of the 3 influenza vaccine strains. These results suggest that a combination of a low dose allosteric mTOR inhibitor (RAD001) plus a catalytic mTOR inhibitor (BEZ235) resulted in greater improvement in influenza vaccination response than RAD001 or BEZ235 monotherapy.

**Fig. 2.**
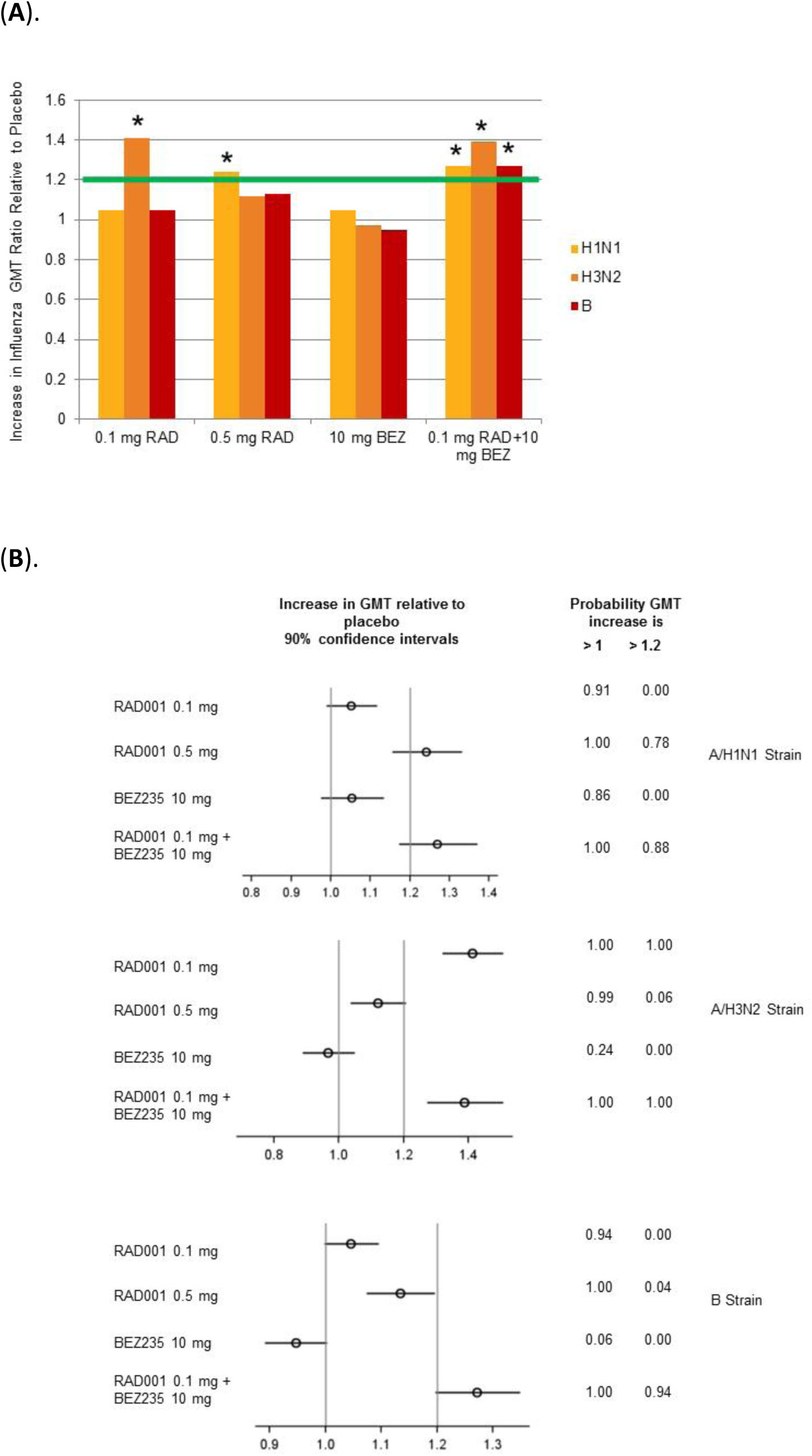
Increase in antibody titers to influenza vaccine strains in mTOR inhibitor treatment groups relative to the placebo treatment group. **A)** Shown is the increase in the ratio (4 weeks after vaccination/baseline) in geometric mean titers (GMT) to each of the 3 influenza vaccine strains (A/H1N1 [A/California/7/2009], A/H3N2 [A/Texas/50/2012], or B [B/Massachusetts/2/2012]) in RAD001 and/or BEZ235-treated cohorts relative to the placebo cohort. The green line indicates the 20% increase in GMT ratios relative to placebo that is required in 2 out of 3 influenza vaccine strains in order to meet the primary endpoint of the study. Asterisks indicate that the probability that the increase in GMT ratio relative to placebo exceeds 1.0 is 100%. **B)** Forest plots of the data presented in (A) including 90% confidence intervals and probability that the GMT ratio as compared to placebo is >1 or > 1.2.

Combinations of low doses of catalytic and allosteric mTOR inhibitors have been reported to inhibit synergistically multiple nodes downstream of TORC1 without inhibiting TORC2 (*14*). Although currently available assays are not sufficiently sensitive to measure TORC1 activity in human peripheral blood samples, concentrations of RAD001 and BEZ235 achieved in the blood of elderly subjects treated with 0.1 mg RAD001 (approximately 0.8 nM Cmax) and 10 mg BEZ235 (approximately 14 nM Cmax) in the current trial have been shown to synergistically inhibit the phosphorylation of both S6K and 4EBP1 downstream of TORC1 *ex vivo* (*15*). To confirm that the combination of low dose RAD001 + BEZ235 is associated with more complete TORC1 inhibition than low dose RAD001 or BEZ235 monotherapy, phosphorylation of the TORC1 substrates S6 kinase, S6 and 4EBP1 was measured in the livers of rats treated for 7 days with the dose equivalent of 0.1 mg RAD001, 0.5 mg RAD001, 10 mg BEZ235 or a combination of 0.1 mg RAD001 and 10 mg BEZ235 (Fig. 3). All mTOR inhibitor dosing regimens except the dose equivalent of 0. 1 mg RAD001 significantly inhibited the phosphorylation of S6K and/or S6. (Fig. 3). However only BEZ235 alone or in combination with RAD001 inhibited the phosphorylation of 4EBP1 (Fig. 3B). Moreover, the only mTOR inhibitor dosing regimen that significantly inhibited all 3 nodes downstream of TORC1 was the combination of RAD001 and BEZ235 (Fig 3B). These data suggest that treatment of elderly subjects with a combination of low dose RAD001 + BEZ235 may result in greater improvement in influenza vaccination response than treatment with RAD001 or BEZ235 monotherapy due to more complete TORC1 inhibition achieved with the combination.

**Fig 3.**
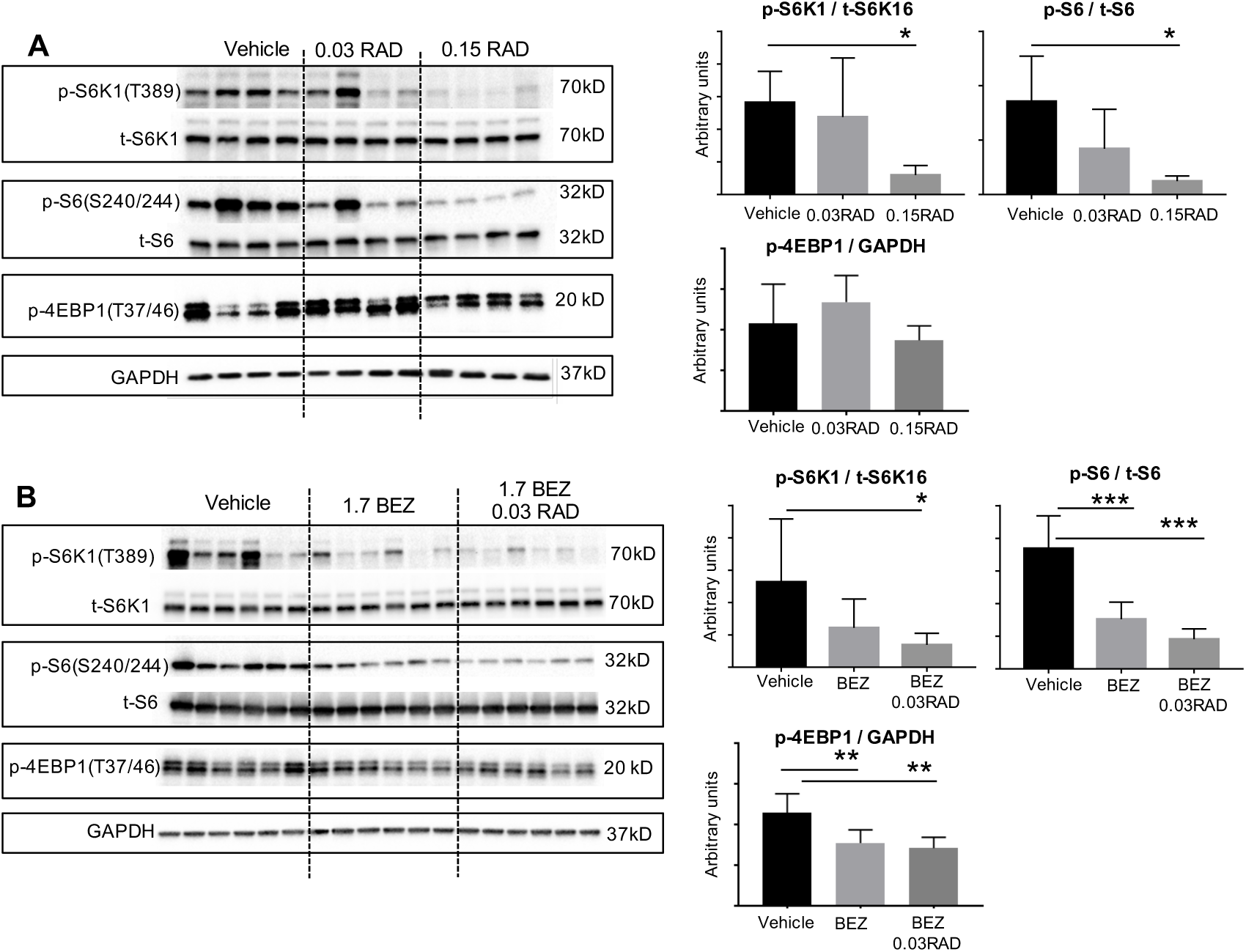
TORCI inhibition associated with low dose mTOR inhibitor regimens. Phosphorylated (p-) and total (t-) protein amounts for S6K1, S6 and 4EBP1 in the livers of rats treated daily for 7 days with RAD001 at the dose equivalent of 0.1 mg in humans (0.03 mg/kg) or 0.5 mg in humans (0.15 mg/kg) (**A**); or BEZ235 given at the dose equivalent of 10 mg in humans (1.7 mg/kg) alone or in combination with the dose equivalent of RAD001 0.1 mg (0.03 mg/kg) (**B**). Tissues were collected 4 hours following the last dose. Each lane in the immunoblots on the left represents a single rat. GAPDH is shown as a loading control. On the right, p-S6K1(T389) and p-S6(Ser240/244) amounts on the immunoblot were quantified relative to their respective total protein amounts by densitometry. Levels of p-4EBP1 (T37/46) were quantified relative to GAPDH protein amounts because we were unable to validate the antibody to total 4EBP1 for rat tissue. Y-axes represent arbitrary units. For each group n = 4-6 rats. Data are mean ± *SD.* Data were analyzed with a one way ANOVA followed by Dunnett’s multiple comparison tests, where means from all groups were compared to the vehicle treated group. * P<0.05; *** P < 0.001. In each figure, “RAD” represents RAD001 and “BEZ” represents BEZ235. The numbers 0.03, 0.15 and 1.7 represent the dose of the corresponding compound in mg/kg/day.

As an additional assessment of immune function, the overall infection rate in each treatment group was assessed by having subjects record any infections they experienced during the year following the initiation of study drug treatment in a diary that was reviewed by study investigators at each study visit. In addition, infections were captured through phone calls between site staff and subjects that that occurred weekly during the 6 weeks subjects were on study drug, and then monthly for the remainder of the trial. The largest and most statistically significant decrease (p=0.001 vs placebo) in the fitted annualized rate of infections reported by subjects was in the RAD001 + BEZ235 combination treatment group (1.49 infections/per person per year (py), 95% confidence interval 1.19-1.86) as compared to placebo (2.41 infections/py, 95% confidence interval 2.00-2.90). The BEZ235 monotherapy treatment group also had a statistically significant (p = 0.008 vs placebo) reduction in the annualized rate of infections reported by subjects (1.61 infections/py, 95% confidence interval 1.28-2.03) (Fig. 4A). There was a trend toward a reduction in infection rates in both RAD001 monotherapy treatment groups but the reductions were not statistically significant.

**Fig. 4.**
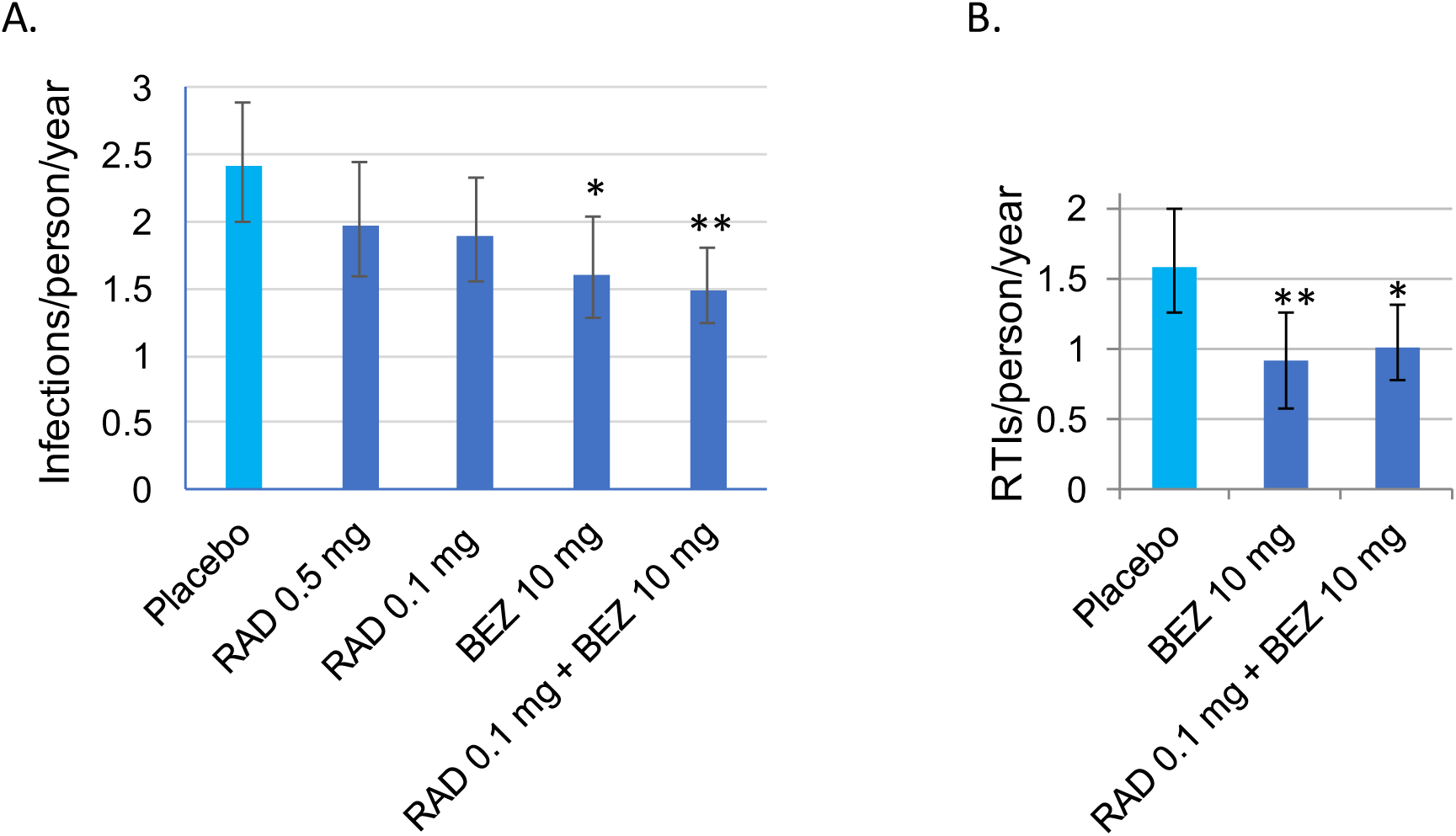
TORC1 inhibition decreases infection rates in the elderly. (**A**). Fitted annual rates of infections reported per person per year in each treatment group are shown. * p=0.008; ** p=0.001 vs placebo. (**B**). Fitted annual rates of respiratory tract infections (RTIs) reported per person per year in the placebo, BEZ235 monotherapy and BEZ235+RAD001 combination treatment groups are shown. * p=0.01, ** p=0.008 vs placebo. In both figures, error bars indicate 95% confidence intervals.

The majority of infections reported during the trial were respiratory tract infections. To determine if a reduction in respiratory tract infections contributed to the significant reduction in total infections reported in the BEZ235 and RAD001+BEZ235 treatment groups, the annualized rate of respiratory tract infections reported in these 2 treatment groups relative to placebo was assessed as a post-hoc analysis. Both BEZ235 monotherapy and BEZ235+RAD001 combination therapy were associated with a significant reduction as compared to placebo in the annualized rate of respiratory tract infections reported by subjects (Figure 4B).

Since the combination of low dose RAD001+BEZ235 was the only mTOR inhibitor dosing regimen associated both with a significant decrease in infections and a significant improvement in the response to 3/3 influenza vaccine strains, subsequent mechanistic studies focused on this treatment group. To explore whether the combination of low dose RAD001+BEZ235 enhanced immune function by decreasing systemic inflammation, several inflammatory cytokines were measured in serum obtained from subjects at baseline and after 6 weeks of either placebo or RAD001+BEZ235 treatment. There were no significant differences in serum levels of interleukin 6 (IL6), interferon gamma (IFN_γ_), tumor necrosis factor alpha (TNFα) or interleukin 18 (IL18) in the RAD001+BEZ235 as compared to the placebo treatment groups (Fig. S1).

To assess other possible molecular mechanisms underlying the enhanced immune function of the RAD001+BEZ235 combination, we conducted mRNA sequencing analysis of whole blood from subjects at baseline and after 6 weeks of either placebo or RAD001+BEZ235 treatment. Whole-blood gene expression data revealed a highly statistically significant, low level up-regulation of pathways related to interferon signaling (Table 2 and Fig. S2). Some of the genes whose expression was most highly upregulated in the enriched pathways were a subset of Type 1 interferon-induced genes that play a critical role in the immune response to viruses (Table 2) (*18*). A complete list of significantly enriched pathways after RAD001+BEZ235 treatment is provided in Table S2. These findings raise the possibility that upregulation of a subset of type 1 interferon-induced genes by a low dose combination of RAD001+BEZ235 may contribute to enhanced immune function and reduced infection rates in the elderly. In particular, the upregulation of antiviral gene expression is likely to underlie the significant reduction in respiratory tract infections seen in the RAD001+BEZ235 treatment cohort since most respiratory tract infections, particularly upper respiratory tract infections, are viral in origin.

**Table 2.** Pathways and genes upregulated after RAD001+BEZ235 treatment as determined by whole blood gene expression analysis

| Pathway | Mean FC<br>Genes In<br>Pathway | p-value | Up-regulated<br>Genes* |
| --- | --- | --- | --- |
| INTERFERON_ALPHA_BETA_SIGNALING | 0.08 | 1e-21.8 | IFI27; IFIT2; IFIT1;<br>IFIT3; MX1;<br>OAS3;<br>ISG15 |
| INTERFERON_SIGNALING | 0.04 | 1e-36.7 | IFI27; IFIT2; IFIT1;<br>IFIT3; MX1;<br>OAS3;<br>HERC5; ISG15 |
| CYTOKINE_SIGNALING_IN_IMMUNE_SYSTEM | 0.02 | 1e-43.5 | IFI27; IFIT2; IFIT1;<br>IFIT3; MX1;<br>OAS3;<br>HERC5; ISG15 |

\* Listed up-regulated genes are those determined to be outliers by the Tukey method of outlier detection.

In conclusion, the current study demonstrated that a combination of low dose RAD001+ BEZ235 led to a significant improvement in immune function as assessed by both the annualized rate of infections reported by subjects and the humoral response to influenza vaccination. Thus our findings suggest that synergistic inhibition of multiple nodes downstream of TORC1 with a combination of low dose RAD001 and BEZ235 results in more significant enhancement in immune function in elderly subjects than low dose RAD001 or BEZ235 monotherapy that lead to less complete inhibition of TORC1.

The results of the current study suggest that the pharmacodynamics effects of mTOR inhibitors are dose-dependent. The doses of RAD001 used in this study are 5-100 fold lower than the doses of RAD001 approved for use in organ transplant and oncology patients. Similarly the dose of BEZ235 used in the current study is 120-fold lower than the maximum tolerated dose established in oncology patients. At higher doses, mTOR inhibitors suppress T cell proliferation and have been associated with increased rates of infection. In contrast, results of this study suggest that very low doses of mTOR inhibitors enhance immune function and decrease infection rates.

The mechanism underlying the improvement in immune function in elderly subjects treated with low doses of mTOR inhibitors is likely to be multifactorial. In a previous study in elderly subjects, we demonstrated that mTOR inhibition with RAD001 monotherapy decreased the percentage of exhausted PD1+ T cells that have defective response to antigens (*16*). Here we demonstrate that the combination of RAD001+BEZ235 also upregulated a subset of interferon-stimulated genes (ISGs) that play a critical role in the innate immune response to pathogens, particularly viruses. There was no change in IFNγ levels in serum after RAD001+BEZ235 therapy (Fig. S1) suggesting that RAD001+BEZ235 may be upregulating signaling pathways downstream of interferons. One possibility is that TORC1 inhibition by low dose RAD001+BEZ235 decreases lipid and cholesterol synthesis within cells due to decreased activation of SREBP2 (*6*). Decreased cholesterol biosynthesis after SREBP2 knockdown has been shown previously to increase expression of a subset of antiviral ISGs (similar to the ISGs upregulated after RAD001+BEZ235 treatment) and protect against viral infection (*19*). The magnitude of ISG upregulation in whole blood after RAD001+BEZ235 treatment was small (an average increase of 17.8% for genes defined as up-regulated in Table 2). A low level increase in ISG expression may be sufficient to enhance immune function in the elderly while avoiding the undesirable adverse events that occur in patients treated with recombinant interferon, in whom much higher levels of ISG induction are observed (*20*).

Surprisingly, both BEZ235 and the combination of RAD001+BEZ235 led to a significant reduction in infections for a year despite the fact that study drug was discontinued after 6 weeks of treatment. These results suggest that mTOR inhibitor therapy may lead to persistent improvements in immune function after drug discontinuation. Persistent beneficial effects of short courses of mTOR inhibitor therapy also have been demonstrated in mice. Specifically, 6-12 weeks of the mTOR inhibitor rapamycin have been shown to extend lifespan in elderly mice (*3,21*).

The decrease in infection rates seen in this study are of clinical relevance since infections, particularly respiratory tract infections, are one of the leading causes of death in the elderly (*5,6,22*). Since hundreds of different viral serotypes cause respiratory tract infections, immunotherapy such as TORC1 inhibition that harnesses the ability of the immune system to fight multiple viral pathogens may be more effective than therapies targeting each individual pathogen. Moreover, the immune system is necessary not only to fight infectious pathogens but also for cancer immunosurveillance and to clear senescent cells that may cause organ damage during aging. Therefore therapies such as BEZ235+RAD001 that enhance immune function may have pleiotropic health benefits in the elderly.

## Acknowledgments

We thank the volunteers, investigators and site staff who participated in this study including Simon Carson, Paul Noonan, Dean Millar-Coote, Edward Watson, Phillipa Murray, Jane Kerr, Jason Prycke, Russell Scott, Paul Dawkins, and Barney Mongomery. We would also like to thank the staff of Pharmaceutical Solutions, particularly Elaine Gent. We thank Walter Carbone and Judith Knehr for the technical performance of the RNA-sequencing experiment and Sharon Wang for performing immunoblotting.

## Funding

Supported by Novartis Institutes of Biomedical Research.

